# The evolution of the natural killer complex; a comparison between mammals using new high-quality genome assemblies and targeted annotation

**DOI:** 10.1101/069922

**Authors:** John C. Schwartz, Mark S. Gibson, Dorothea Heimeier, Sergey Koren, Adam M. Phillippy, Derek M. Bickhart, Timothy P. L. Smith, Juan F. Medrano, John A. Hammond

**Author notes:** Current: CEDOC, Faculdade de Ciências Médicas, Universidade Nova de Lisboa, 1150-082 Lisbon, Portugal (Mark S. Gibson).

## Abstract

Natural killer (NK) cells are a diverse population of lymphocytes with a range of biological roles including essential immune functions. NK cell diversity is created by the differential expression of cell surface receptors which modulate activation and function, including multiple subfamilies of C-type lectin receptors encoded within the NK gene complex (NKC). Little is known about the gene content of the NKC beyond rodent and primate lineages, other than it appears to be extremely variable between mammalian groups. We compared the NKC structure between mammalian species using new high quality draft genome assemblies for cattle and goat, re-annotated sheep, pig and horse genome assemblies and the published human, rat and mouse lemur NKC. The major NKC genes are largely in syntenic positions in all eight species, with significant independent expansions and deletions between species, allowing us to present a model for NKC evolution during mammalian radiation. The ruminant species, cattle and goats, have independently evolved a second *KLRC* locus flanked by *KLRA* and *KLRJ* and a novel *KLRH*-like gene has acquired an activating tail. This novel gene has duplicated several times within cattle, while other activating receptor genes have been selectively disrupted. Targeted genome enrichment in cattle identified varying levels of allelic polymorphism between these NKC genes concentrated in the predicted extracellular ligand binding domains. This novel recombination and allelic polymorphism is consistent with NKC evolution under balancing selection, suggesting this diversity influences individual immune responses and may impact on differential outcomes of pathogen infection and vaccination.

## Introduction

Natural killer (NK) cells are a diverse population of circulating lymphoid cells with cytotoxic and cytokine-secreting functions, particularly in response to intracellular pathogen infections and neoplasms. Although rare, primary NK cell immunodeficiency leads to complications and/or death from severe herpesviral infections, virus-associated tumor growth, leukemia, and mycobacterial infections (Orange 2013). Dysregulated MHC class I expression on nucleated cells, such as during viral infection, is recognized by a diverse repertoire of NK cell surface receptors which mediate their immune functions through direct recognition of equally diverse MHC class I molecules. In mammals, NK cell receptors for MHC class I are encoded within two unrelated and independently segregating gene complexes; the leukocyte receptor complex (LRC) containing genes encoding the killer cell immunoglobulin-like receptors (*KIR*), and the natural killer complex (NKC) containing multiple members of killer cell lectin-like receptor genes (*KLR*). Both gene complexes evolve rapidly, vary in gene content within and between species and can encode both activating and inhibitory polymorphic receptors. Thus, a highly diverse NK cell repertoire containing multiple highly similar receptors allows for a finely-tuned ability to discriminate MHC class I expression between healthy and damaged cells.

The number of *KIR* and *KLR* genes is highly variable between mammalian species, with primate and rodent species the best studied to date (Guethlein et al. 2015). Humans and other higher primates have an expanded, highly polymorphic and gene variable KIR locus, and possess four functional *KLRC* genes (*NKG2A*, *C*, *E*, and *F*) and a single *KLRA* (*Ly49*) gene or pseudogene (Wende et al. 1999; Wilson et al. 2000; Trowsdale et al. 2001). In contrast, rats (*Rattus norvegicus*) and mice (*Mus musculus*) possess one and two *KIR* genes, respectively, but these are not located in the LRC and are thought to have alternative functions. However, rodents have a highly expanded and diverse repertoire of *KLRA* genes (Anderson et al. 2001; Hoelsbrekken et al. 2003; Higuchi et al. 2010) and a unique, yet related *KLRH* gene (Naper et al. 2002). The primate *KIR* and the rodent *KLRA* and *KLRH* bind classical MHC class I molecules to control NK cell function, a rare example of convergent evolution that illustrates the fundamental importance of this receptor-ligand system.

Beyond rodents and humans, a few other species have been studied in some detail. Horses (*Equus caballus*), for example, possess an expanded *KLRA* repertoire of five polymorphic genes and a single putatively functional *KIR3DL*-like gene (Takahashi et al. 2004; Futas and Horin 2013). The mouse lemur (*Microcebus murinus*) has expanded a different NKC gene family, possessing five functional *KLRC* and three functional *KLRD* (*CD94*) (Averdam et al. 2009). Together, *KLRC* and *KLRD* form a heterodimeric pair providing the mouse lemur with a significantly expanded *KLRC/KLRD* combinatorial repertoire. However, NK receptor diversification is not always a prerequisite for a species survival. Several marine carnivores, (seals and sea lions) possess a single functional *KIR* and single functional *KLRA*, while their terrestrial relatives, cats (*Felis catus*) and dogs (*Canis lupus*), only appear to have a functional *KLRA*, with the *KIR* gene being disrupted or deleted, respectively (Hammond et al. 2009). Pigs (*Sus scrofa*) also possess a single *KIR* and a single *KLRA*, yet it is uncertain if either these genes are functional (Gagnier et al. 2003; Sambrook et al. 2006).

Findings to date indicate that cattle (*Bos taurus*) are unique in having expanded and diversified NK cell receptor genes within both the NKC and LRC (McQueen et al. 2002; Birch and Ellis 2007; Guethlein et al. 2007). Cattle possess at least seven *KLRC*, two *KLRD*, and a single but polymorphic *KLRA* within the NKC (Birch and Ellis 2007; Dobromylskyj et al. 2009), and eight functional *KIR* genes in the LRC (Sanderson et al. 2014). However, the characterisation of the NKC relies largely on the current public genome assemby (Elsik et al. 2009). Immune gene complexes, however, are often highly repetitive and polymorphic, creating significant assembly problems during whole genome sequencing attempts. As a consequence, several draft genome assemblies indicate that the NKC has had a complex and dynamic evolutionary history during mammalian radiation, a hallmark of strong positive selection, but the genome sequence of these regions and associated annotation is either preliminary or lacking.

An accurate NKC genome sequence and correct annotation is essential to inform functional genomic studies. In an age of heightened concern for food security, this is particularly important for immunogenetic variation in food producing species that could be exploited to improve resilience to infectious diseases. To address this we have improved and confirmed the assembly of the cattle and goat (*Capra hircus*) NKC using BAC sequencing and recent long-read genome assemblies (Smith and Medrano, unpublished) (Bickhart et al. 2016). These were newly annotated, as were the available draft reference genomes for the sheep (*Ovis aries*) (The International Sheep Genomics Consortium et al. 2010), pig (Groenen et al. 2012), and horse (Wade et al. 2009). These were then compared to the well characterised NKC structures of the rat (Flornes et al. 2010), human (Hofer et al. 2001), and mouse lemur (Averdam et al. 2009) allowing us to propose a model for the NKC evolution during the past approximately 92 million years. To additionally assess the level of intraspecies variability in cattle and genes under selection, we investigated polymorphism within the NKC of twenty-three individuals including breeds from both *B. taurus* and *B. indicus*, whose wild ancestors began to diverge approximately two million years ago (mya) (Hiendleder et al. 2008) and which were domesticated separately about 10,000 years ago.

## RESULTS

### Re-assembly of the cattle and goat NKC

Highly repetitive gene complexes are notoriously difficult to assemble and annotate during whole genome sequencing attempts. In the current best public cattle genome assembly (UMD_3.1), the NKC region spanning *KLRA* to *KLRE* is a 730 kb scaffold containing 14 gaps (Figure 1a). The region appears largely intact, but there are clear annotation errors and small contigs that are likely erroneous. Recent resequencing efforts using long-read sequencing technology have produced an improved genome assembly (ARS-UCDv0.1) with >50x higher contiguity than the UMD_3.1 or Btau_5.1 public assemblies (Smith and Medrano, unpublished). Contigs containing the NKC region from the ARS-UCDv0.1 assembly were identified via BLASTN, revealing a single ungapped contig of approximately 12.7 mb containing 785 kb between *KLRA* and *KLRE*. Comparison of this contig to the UMD_3.1 scaffold revealed that the overall structure is almost identical; however, within the UMD_3.1 assembly we identified three mis-ordered contigs, a 16 kb contig containing olfactory receptors, and a sizeable sequence gap of approximately 70 kb downstream from *KLRA* (Figure 1a). To confirm which NKC assembly was accurate, we sequenced and assembled seven bacterial artificial chromosome (BAC) clones containing cattle NKC sequence. Five clones were from two Friesian bulls (three from the TPI-4222 library (Di Palma 1999) and two from the RPCI-42 library; http://bacpac.chori.org/) and two clones were from a Hereford bull (CHORI-240 library; http://bacpac.chori.org/). The Hereford BAC library was the source for the minimum BAC tiling path sequenced as the primary basis for the existing UMD_3.1 assembly, supplemented by whole genome shotgun sequence from a daughter (L1 Dominette 01449) of the same bull (Elsik et al. 2009), who in turn was the source of the ARS-UCDv0.1 long-read assembly. BAC assemblies for the TPI clones were confirmed by comparing their restriction digest band sizes to their *in silico* predictions. All seven clones mapped with high identity to both assemblies but no structural differences were identified between the BAC clones and the ARS-UCDv0.1 genome assembly (Figure 1a). This confirmed that the sequence gaps, contig mis-ordering, and placement of putative olfactory receptor genes are errors in UMD_3.1.

**Figure 1.**
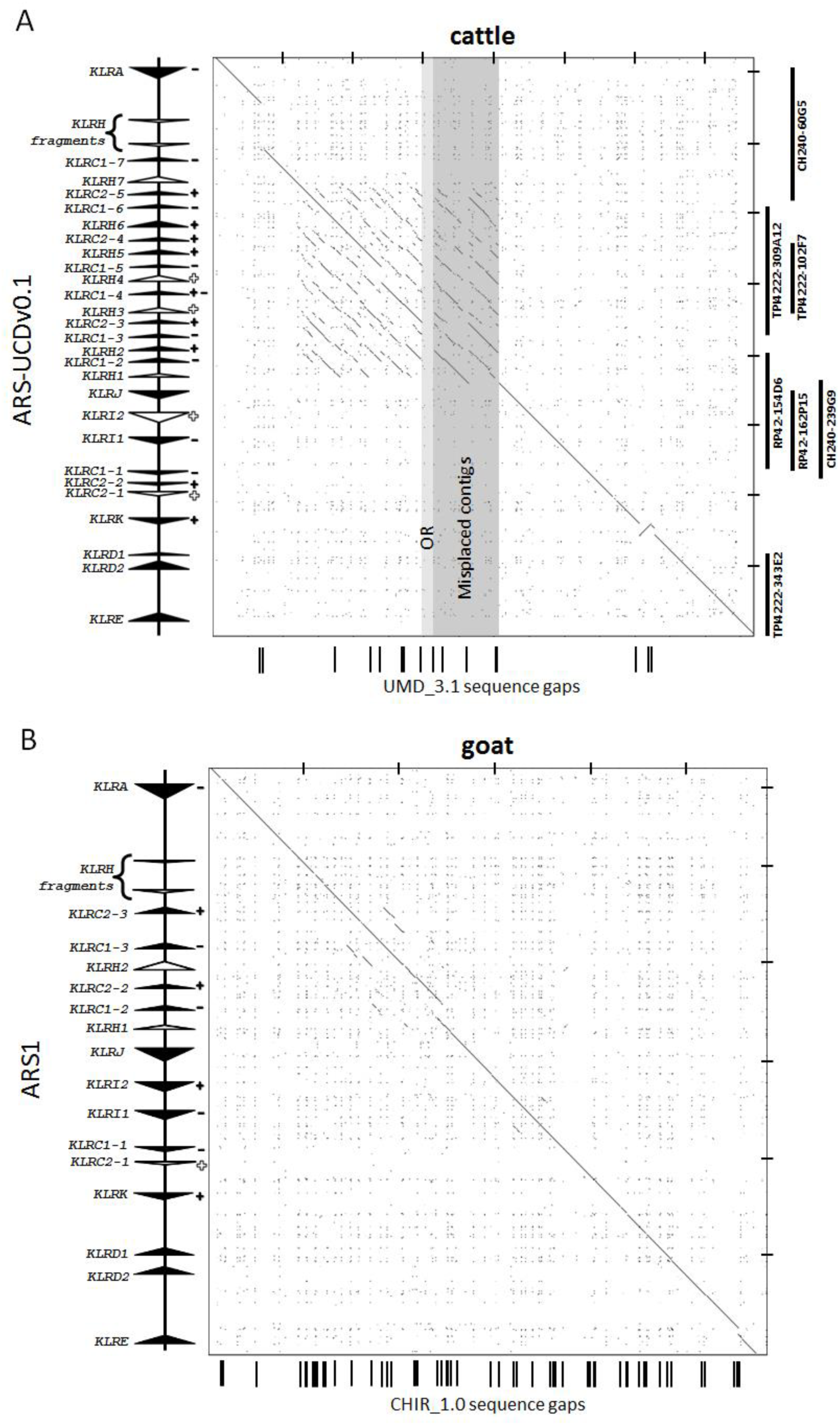
Recurrence plots of cattle (**a**) and goat (**b**) NKC regions comparing the sequence identities of the reference genome assemblies (*x*-*axes*) with the current respective long-read assemblies (*y*-*axes*). Gene annotation is shown at left. Genes which are either putatively functional (*closed arrows*) or non-functional (*open arrows*) are indicated and point in the direction of transcription. Genes which encode receptors that possess inhibitory (−) and/or activating components (+) are indicated, and open symbols denote non-functional genes. Gaps within the reference assemblies are represented by black bars below the *x*-*axes*. No sequence gaps were present within either long-read assembly. Tic-marks at *top* and *right* are separated by 100 kb. Misplaced and olfactory receptor (*OR*)-containing contigs are indicated for the cattle genome as grey boxes. BAC clones used in the current analyses are represented at *right*.

We performed a similar analysis with the current public goat genome assembly (CHIR_1.0) and a new long-read *de novo* goat assembly, ARS1, that has also used PacBio sequencing to improve assembly contiguity (Bickhart et al. 2016). In the CHIR_1.0 assembly, the NKC region from *KLRA* to *KLRE* spans approximately 584 kb and contains a total of 53 sequence gaps (Figure 1b). In contrast, a single contiguous region of approximately 600 kb was identified on a scaffold of approximately 113 mb in the ARS1 assembly. Although these regions are structurally very similar, the long-read assembly resolved the numerous gaps in CHIR_1.0 and included an extra 15 kb of sequence containing a unique NKC gene (*KLRC1-2*, Figure 1b). Our findings with the cattle long-read assembly provide high confidence that the goat ARS1 assembly is accurate, and the scaffolds have been verified by both optical map and chromatin conformation analysis (Bickhart et al. 2016). Therefore we did not repeat the BAC-based analysis that was done for cattle. The complete cattle and goat NKC from the long-read assemblies were manually annotated at high resolution to identify all of the exons related to NKC genes and examine which had the potential to encode functional genes.

### The unique organization and gene expansions within the ruminant NKC

We compared our cattle and goat assemblies to the well characterized human and rat NKC (Hofer et al. 2001; Flornes et al. 2010) to examine the evolution of the ruminant NKC. The general organization of the NKC is largely conserved across the four species, with species-specific expansions and contractions within relatively defined zones. Notably, the human NKC is relatively compact and encodes only six functional genes; *KLRK*, *KLRD*, and four copies of *KLRC* (Figure 2), and lacks *KLRI* and *KLRE* genes. Multiple copies of *KLRC* are also encoded between *KLRA* and *KLRK* in all four species, whilst *KLRA* is always centromeric to *KLRC*. *KLRJ*, a gene most closely related to *KLRA*, is localized telomeric from *KLRA* and centromeric from the *KLRI* genes in both the cattle and goat genomes. Encoding both a transmembrane (TM) and a cytoplasmic tail, *KLRJ* lacks either activating or inhibitory components making its function ambiguous, suggestive of a heterodimeric role as seen with *KLRD* and *KLRE*, albeit with an unknown partner.

**Figure 2.**
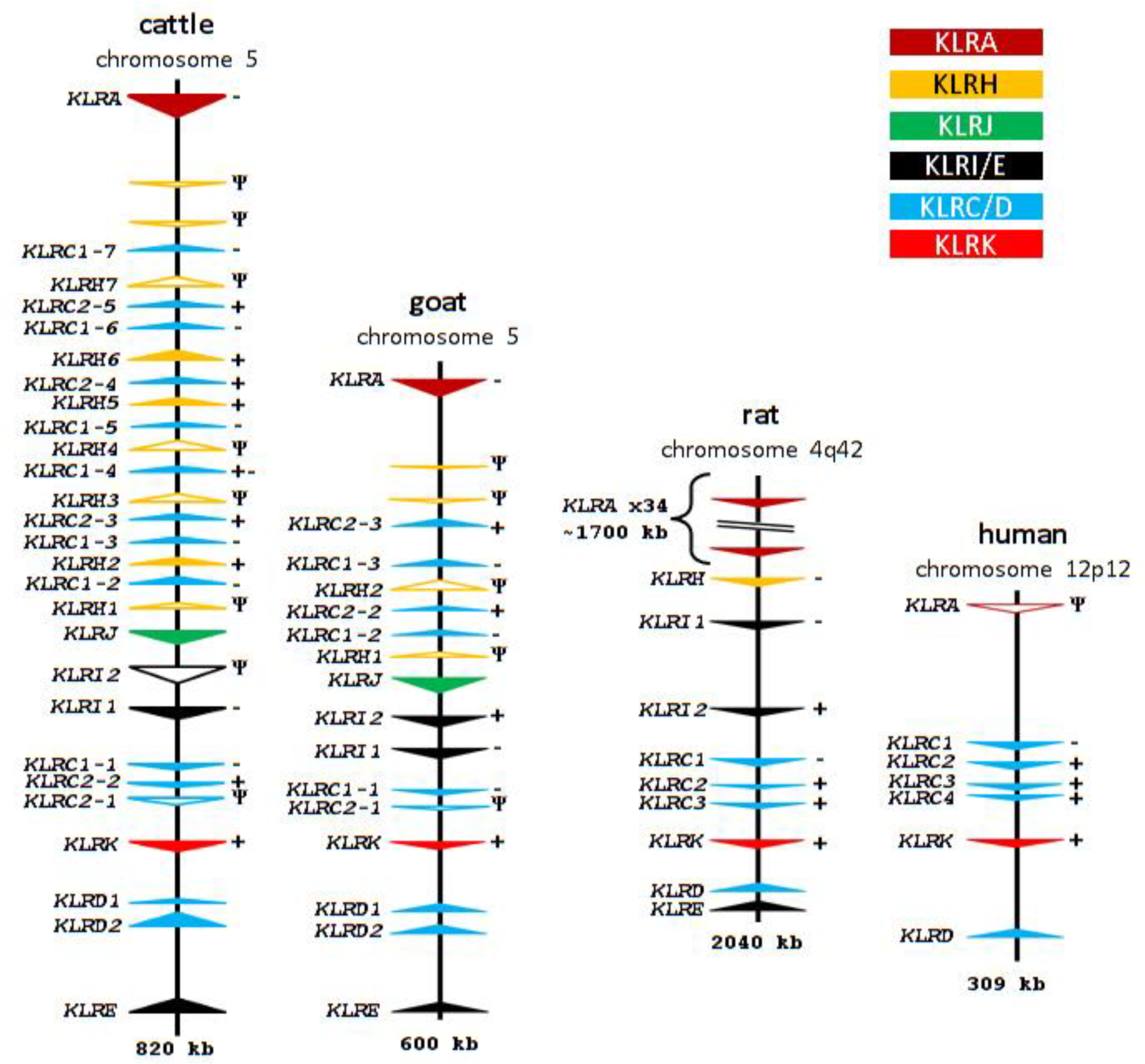
Comparative organization of the NKC in selected species. Genomic regions are approximately to scale, visualized as in Figure 1 with *Ψ* indicating pseudogenes, and anchored on *KLRK*. Rat and human NKC organizations are adapted from Flornes et al. (2010) and Hofer et al. (2001), respectively.

Immediately downstream from *KLRA* in both cattle and goats are two ~150 bp vestigial *KLRH*-like exons (approximately 72% sequence identity with rat *KLRH1*). Recurrence plot sequence identity analysis revealed the presence of a highly repetitive region approximately 280 kb in size midway between *KLRA* and *KLRJ* (Figure 1a). Most interestingly, this region in both cattle and goats contains a novel, expanded assortment of C-type lectin-like genes encoded in the opposite orientation to *KLRA*. In cattle, this includes 16 novel genes, nine of which are *KLRC*-like (Figure 3a and b), interspersed with seven genes most closely related to rat *KLRH1* by phylogenetic analysis of their extracellular C-type lectin domain (Figure 3b and Supplementary Figure 1). Five of these seven *KLRH*-like genes bear activating *KLRC2*-like cytoplasmic and TM domains (Figure 3a). This region in goats has likewise expanded to include four *KLRC* genes and two *KLRH*-like genes (Figure 2), one of which possesses exons 1 and 3 of a *KLRC2*-like activating tail. It is therefore apparent that *KLRC* expansion into this region as well as recombination with *KLRH* preceded the Bovinae-Caprinae divergence ~30 mya (Hiendleder et al. 1998).

**Figure 3.**
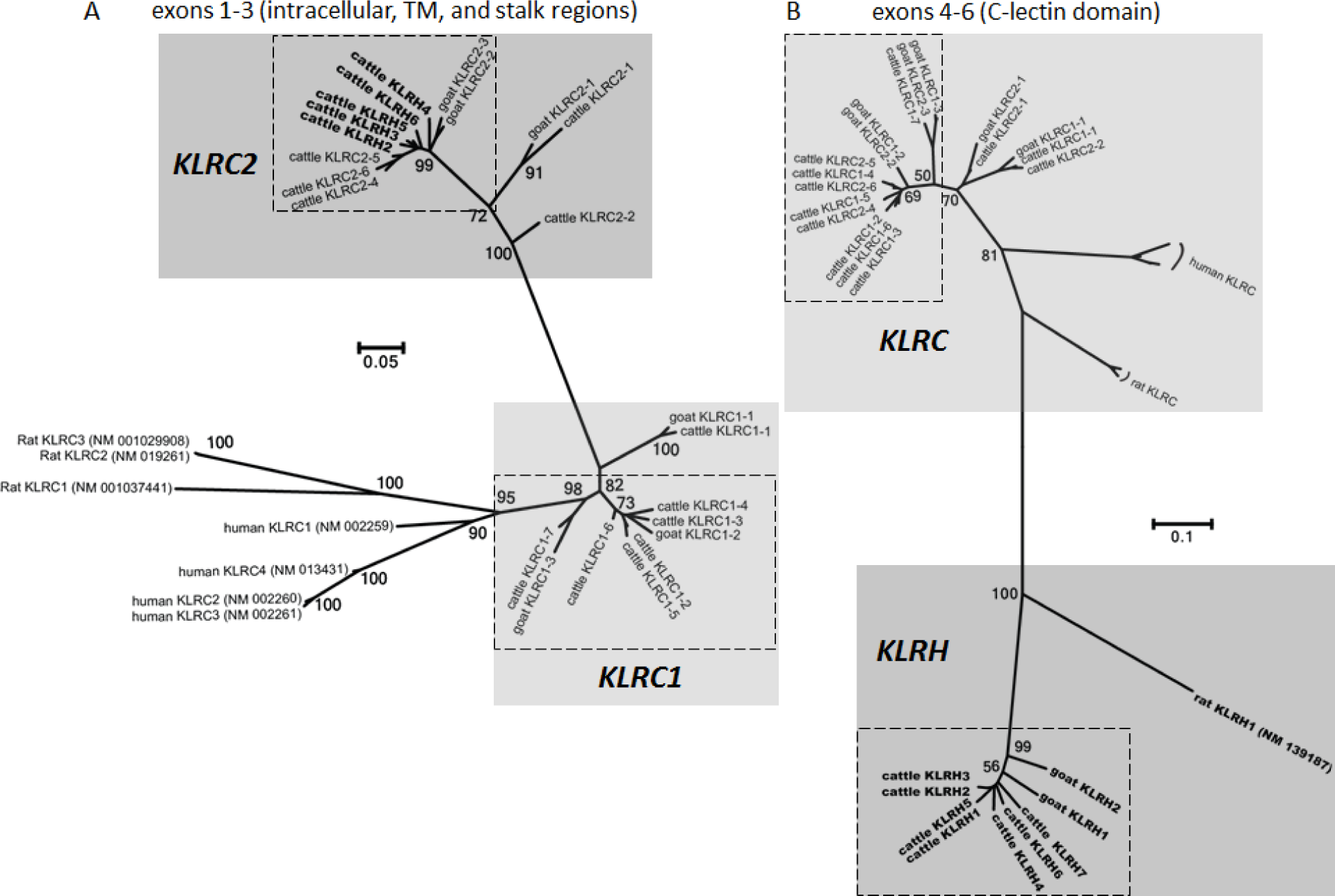
Phylogenetic relationships of coding region sequence for *KLRC* and *KLRH* cytoplasmic and TM regions (**a**) and lectin domain in cattle, goats, humans and rats (**b**). Nucleotide sequence alignments were generated using ClustalW (Thompson et al. 1994) and the trees were constructed using maximum likelihood based on the Tamura-3 parameter (Tamura 1992) model within MEGA6 (Tamura et al. 2013) using 100 bootstrap repetitions. Nodes supported by >50 bootstraps are labelled. The first three exons of rat *KLRH* were excluded from (**a**) as the sequence was too divergent to be aligned. *Dashed boxes* indicate ruminant genes found within the expanded region flanked by *KLRA* and *KLRJ*. To ease visualization, *KLRH* genes are shown in ***bold***.

Five of the six *KLRC1* genes in this expanded region of cattle are >95% identical to one another, indicative of more recent evolutionary expansion. All six inhibitory *KLRC1* genes and all three activating *KLRC2* genes are putatively functional, based on their open reading frame sequences and the conservation of canonical splice site motifs. An unusual feature was previously reported in which a cattle *KLRC1* cDNA (*NKG2A-07*) possessed a cytoplasmic tail containing two immunoreceptor tyrosine-based inhibition motifs (ITIMs) and a predicted TM region containing an arginine residue creating the potential for both inhibitory and activating functions, respectively (Birch and Ellis 2007). The existence of this gene was confirmed in the ARS-UCDv0.1 genome assembly, as it matches to *bota_KLRC1-4* which is located in the center of the expanded cattle *KLRC* region (Figure 2).

The other cluster of *KLRC* genes flanked by *KLRK* and *KLRI* in rat, goat, and cattle, appears more conserved across these species. Both the cattle and goat genomes contain a single inhibitory *KLRC1* gene and a *KLRC2* pseudogene, whilst cattle possess an additional likely functional *KLRC2* (Figure 2). The *KLRC2* pseudogenes of cattle and goats share the same disabling features and are both missing the last two exons indicating that functionality was lost prior to their divergence from a common ruminant ancestor. Importantly, *KLRD* has duplicated in both cattle and goats, consistent with an expanded *KLRC* repertoire in other species and suggesting that the heterodimeric *KLRC/KLRD* partnership has been preserved and subject to similar diversification pressures.

### *KLRH* has been reactivated and expanded in cattle

The existence of *KLRH*-like genes in ruminants is intriguing, as *KLRH* has not yet been described beyond the rodent lineage. Five such genes in cattle possess activating TM and cytoplasmic domains (*bota_KLRH2*, *bota_KLRH3*, *bota_KLRH4*, *bota_KLRH5*, and *bota_KLRH6*) of which three are putatively functional (*bota_KLRH2*, *bota_KLRH5*, and *bota_KLRH6*), although one of these contains two non-canonical splice sites (*bota_KLRH6*). Furthermore, one of two *KLRH* genes in the goat has likewise aquired a *KLRC*-like tail (*cahi_KLRH2*). However, exon 2 of this tail appears to have been subsequently deleted from *cahi_KLRH2*. Moreover, two non-canonical splice sites in exons 3 and 4 and a frameshift in exon 5 have likely destroyed its functionality, despite maintenance of an open reading frame across the exon 1 to exon 3 putative fusion. Two additional genes in cattle (*bota_KLRH1* and *bota_KLRH7*) and one in goats (*cahi_KLRH1*) do not appear to be associated with exons encoding the N-terminal intracellular and TM domains. An 80 bp fragment most similar to a *KLRA*-like N-terminal region was identified approximately 4 kb upstream of *cahi_KLRH1* using the National Center for Biotechnology Information (NCBI) conserved domain database (Marchler-Bauer et al. 2015), whereas BLASTN, BLASTX, and HMMgene failed to predict this portion of the gene. Despite this, both *cahi_KLRH1* and *bota_KLRH1* intriguingly form an intact open reading frame across the exons encoding the C-terminal lectin domain. Together, these findings indicate that *KLRH* was functionally resurrected with the acquisition of an activating tail prior to the Bovinae-Caprinae divergence, and further expanded in the bovine lineage.

### Receptors with activating potential have been disrupted

All four of the cattle pseudogenes and both goat pseudogenes that contain the full complement of exons (*bota_KLRH3*, *bota_KLRH4*, *bota_KLRC2-1*, *cahi_KLRH2*, *cahi_KLRC2-1*) encode potentially activating receptors (Figure 2). It is interesting to note that none of the genes share the same disrupting features. Whereas *bota_KLRH4* contains a single frameshift in exon 5, *bota_KLRH3* contains no disabling genetic lesions within the lectin domain-encoding exons, but contains a frameshift within exon 3 and non-canonical splice site in exon 2. The cattle *bota_KLRH7* gene is disabled by nonsense mutations in both exons 4 and 5 and a non-canonical splice site and frameshift in exon 5, while the other *KLRH*-like gene lacking a tail (*bota_KLRH1*) possesses an intact open reading frame with canonical splice sites. We conclude from this diversity of disabling mutations that both the recombinant *KLRH* genes with activating *KLRC2*-like tails and those apparently lacking tails were independently disrupted after their expansion.

The initiation codon of *bota_KLRI2* is mutated (ATG->AAG) in both the UMD_3.1 and ARS-UCDv0.1 assemblies. Although a potential alternative start site exists several codons downstream, a non-canonical splice site at the end of exon 1 may additionally disrupt the gene. We observed an additional 2 bp frameshift that results in multiple downstream stop codons in both of the overlapping RPCI-42 BAC clones, arguing that the gene has become non-functional in the cattle genome. In contrast, no disabling mutations were observed within the genes encoding inhibitory receptors. This type of selective disruption of activating genes is also a feature of the cattle *KIR* (Sanderson et al. 2014), as well as human *KIR* and mouse *KLRA* (Abi-Rached and Parham 2005).

### Allelic polymorphism is concentrated in the predicted extracellular and ligand binding domains

Allelic polymorphism is a feature of expanded NK cell receptor complexes (Trowsdale et al. 2001). We used the fact that the cattle BAC clones were derived from three different individuals to examine the allelic variability between the overlapping sequences. No SNP variation was identified between the Friesian-derived TPI4222-102F7 and TPI4222-309A12 clones, suggesting they were derived from the same haplotype, which may be a consequence of the historically or current low effective population size for this breed. More surprisingly, there was little polymorphism between these clones and the ARS-UCDv0.1 assembly, despite the latter being derived from a Hereford. Across 180 kb of overlap between the assembly and TPI4222-309A12, there were a total of 177 SNPs and 78 missing nucleotide positions. Only one of these was located within an exon, a synonymous SNP within exon 2 of *bota_KLRH3*. Similarly none of the 713 SNPs between TPI4222-343E2 and the genome assembly were within the coding regions of *bota_KLRE* or *bota_KLRD2*. The CH0RI-240 clones, which were used as a major part of the UMD_3.1 assembly and were derived from L1 Domino 99375, the sire of the animal used for the long-read assembly, contained no SNPs relative to the ARS-UCDv0.1 assembly, perhaps because they came from the same allele assembled from his daughter's genome. However, the CH240-239G9 and RP42-154D6 sequences share a 3 bp insertion within one of the lectin domain-encoding exons of *bota_KLRJ*. Three additional non-synonymous changes were further observed in the lectin region of *bota_KLRJ* on RP42-162P15, but not on RP42-154D6, relative to the genome assembly. Thus, both overlapping RPCI-42 clones derive from different haplotypes. Despite this, both of these clones possess a shared, identical copy of *bota_KLRI2* indicating that these two haplotypes have either recombined or that the *bota_KLRI2* paralog common to both has undergone gene conversion.

The identification of allelic variability in the BAC clones and genome motivated further investigation of polymorphisms within the NKC. We therefore enriched, sequenced and mapped NKC genomic DNA representing husbanded and feral cattle from twenty *B. taurus* and three *B. indicus* (estimated divergence time, 1.7 - 2.0 mya; Hiendleder et al. (2008)). Accurate detection of polymorphism in the expanded 300 kb *KLRC/H* region was not practical using these short reads, due to the highly similar and repetitive nature of these genes and pseudogenes that complicates accurate read mapping. However, high-confidence mapping was possible outside of this region which revealed substantial allelic variation among the remainder of the NKC genes. In total, 77 SNPs (55 non-synonymous) were identified within the coding regions of *KLRA*, *KLRJ*, *KLRI2*, *KLRI1*, *KLRK*, *KLRD1*, *KLRD2*, and *KLRE* (Figure 4). Notably, we observed no sequence variation within the lectin domain of *KLRK* across all 23 animals. Similarly, *KLRD2* and *KLRE* were almost monomorphic, the exception being an apparent divergent *KLRE* genotype observed in all three *B. indicus* animals. In contrast, 14 non-synomymous SNPs were identified within the coding regions for *KLRA* and 11 in *KLRD1*. The former of which confirms a previous report identifying two divergent *KLRA* allelic lineages in cattle (Dobromylskyj et al. 2009). For each of these two genes there appeared to be two major haplogroups, suggesting they may have distinct functional properties. There was no clear relationship between *KLRD1* or *KLRA* genotypes and cattle breed as individuals from different breeds within both *B. taurus* and *B. indicus* share almost identical alleles (Figure 4), which suggests diversifying selection has been operating on these genes. Indeed, highly significant evidence for diversifying selection within the *KLRD* locus of prosimians (but not simians) has been reported based on differential synonymous to non-synonymous substitution rates (Averdam et al. 2009). Furthermore, as our probes captured the flanking genes *STYK1* and *MAGOHB*, we assessed heterozygosity in the flanking region upstream from *KLRA*. Although the diversity of these genes is somewhat limited compared to those of the NKC, their heterozygosity largely corresponds to that observed across the NKC for the 23 animals we assessed (Figure 4).

**Figure 4.**
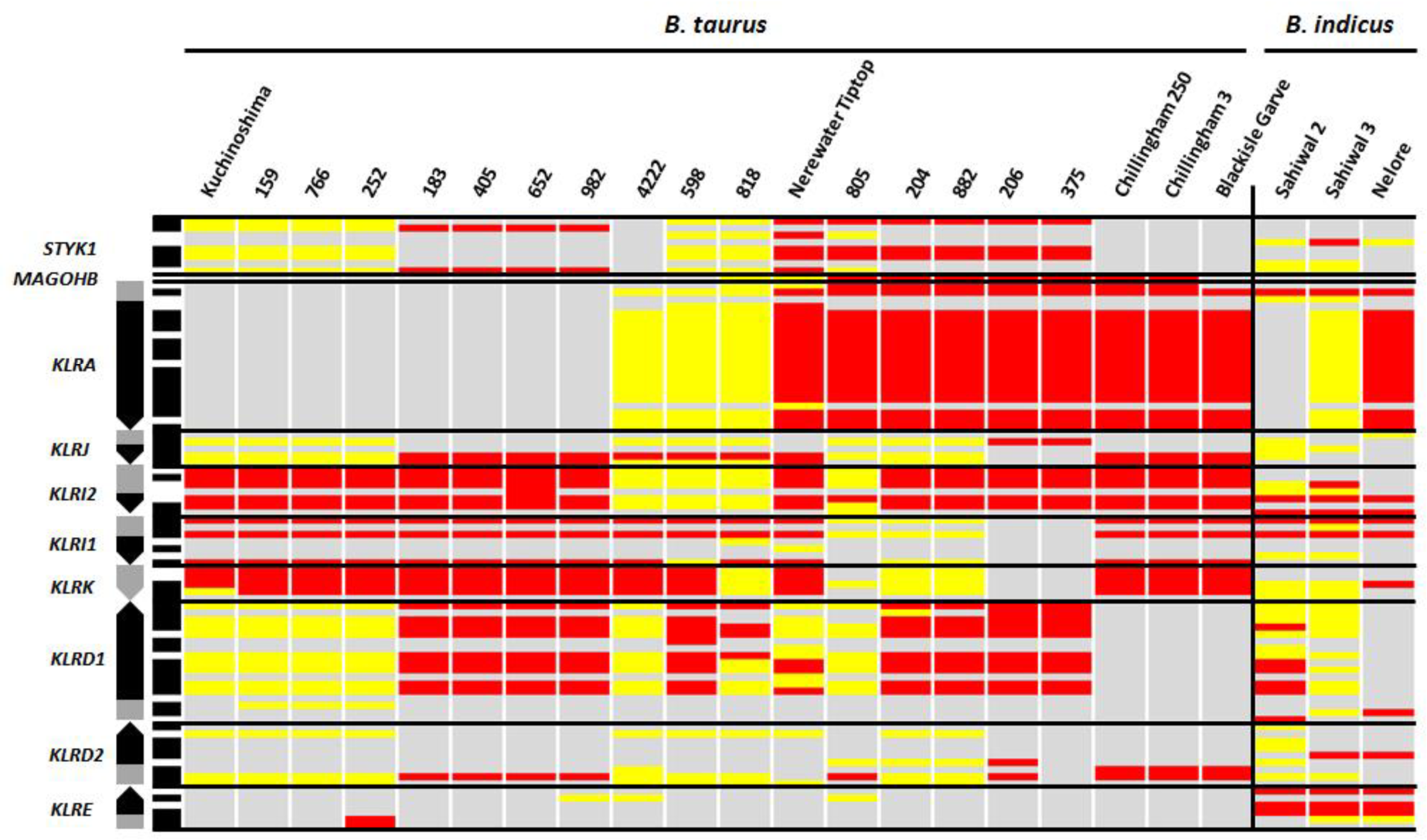
Genetic variation within the cattle NKC coding regions. Genomic orientation is preserved with gene orientation shown at left with arrows pointing in the direction of transcription. *Black-shaded* regions of genes indicate the lectin-coding domains and *grey-shaded* regions indicate cytoplasmic, TM, and stalk regions. Shaded bars at left indicate whether SNPs are synonymous (*white*) or non-synonymous (*black*) when compared to the reference genome (UMD_3.1). *Red-colored* bars indicate homozygous SNPs (approx. 100% of reads), *yellow-colored* bars indicate heterozygous SNPs (approx. 50% of reads), and grey-colored bars indicate identity to the reference.

### Comparison to other mammalian genome drafts confirms the plasticity of NKC genes and the unique organisation in ruminants

To better understand the history of NKC evolution we compared the cattle and goat long read assemblies with the available reference genomes for sheep (0ar_v3.1), pig (Sscrofa10.2), and horse (Equ Cab 2), that are all based on short sequence reads. As expected based on our findings with the UMD_3.1 and CHIR1.0 assemblies, the NKC of these reference genomes contain numerous sequence gaps, making conclusions about detailed genomic structure and allele content provisional (Supplementary Figure 2). However, the sheep NKC assembly is consistent with the goat and cattle, with the only substantial differences being within the highly variable region containing the expanded *KLRC* and *KLRH* which is heavily fragmented into 17 contigs. Furthermore, while there are numerous *KLRC*-like fragments present on various contigs in the sheep assembly, it is impossible to determine whether any of them are associated with a *KLRH*-like gene, as seen in cattle and goat.

The pig is a more distantly related artiodactyl to ruminants, sharing a common ancestor approximately 60 mya (Meredith et al. 2011). They possess a single inhibitory *KLRA* and *KLRJ* with two small *KLRH*-like fragments proximal to and in the same orientation as *KLRA*, similar to the ruminant genomes. In addition, a single *KLRH* gene lacking the first three exons is proximal to *KLRJ* in a syntenic position and in the same orientation as *KLRH1* in ruminants. Overall, the pig NKC is considerably more compact and shows little evidence of gene expansion (Supplementary Figure 2). Notably, the porcine NKC has only a single activating gene, *KLRK*. A single inhibitory *KLRI1*, a single inhibitory *KLRC1*, and a single gene each of *KLRD* and *KLRE* complete this region of the porcine NKC. Thus, pigs possess the most restricted NK cell repertoire so far studied, having both a compact NKC and single *KIR* gene.

The horse shared a common ancestor with cattle approximately 80 mya (Meredith et al. 2011). In the current genome assembly we identified NKC genes spanning a large region of approximately 1260 kb, which contains 13 sequence assembly gaps. We identified five functional inhibitory *KLRA* genes and a single *KLRA* pseudogene, which is consistent with previous studies based on cDNA evidence (Takahashi et al. 2004). Two *KLRH*-like genes are also present in syntenic positions to those found in the artiodactyl genomes, and likewise lack the first three exons. As in pigs, horses appear to possess a single functional gene each of *KLRJ*, *KLRD*, *KLRE*, inhibitory *KLRI1*, and activating *KLRK*. Intriguingly, the *KLRC* locus flanked by *KLRI1* and *KLRK* appears to be substantially expanded compared to all other known genomes. Seven putatively functional inhibitory *KLRC1* genes, three *KLRC2* genes (of which only two have activating motifs), and three *KLRC1* pseudogenes were identified in total. Thus, despite expansion, the horse *KLRC* gene cluster has retained only two putatively functional activating members.

Due to the presence of *KLRJ* within the genomes of the species we studied, we revisited the well characterised NKC of the human, mouse lemur, and rat. While not present in either the human (GRCh38.p5) or the rat (Rnor_6.0) genomes, *KLRJ* is present in the mouse lemur NKC BAC assembly (GenBank: FP236838, Averdam et al. (2009), Supplementary Figure 2). Interestingly, we also identified two *KLRH* genes in the lemur (but not in the human) in the opposite orientation as *KLRA* and *KLRJ*, as seen in the other species (Supplementary Figure 2). As in the other non-human, non-rodent genomes that we investigated, neither of these *KLRH* genes appear to be associated with coding sequence for intracellular or TM regions. Using the annotation of NKC genes in this study in combination with the divergence times between the common ancestors of each species we are able to propose a model for the expansion and contraction of the NKC during mammalian radiation (Figure 5).

**Figure 5.**
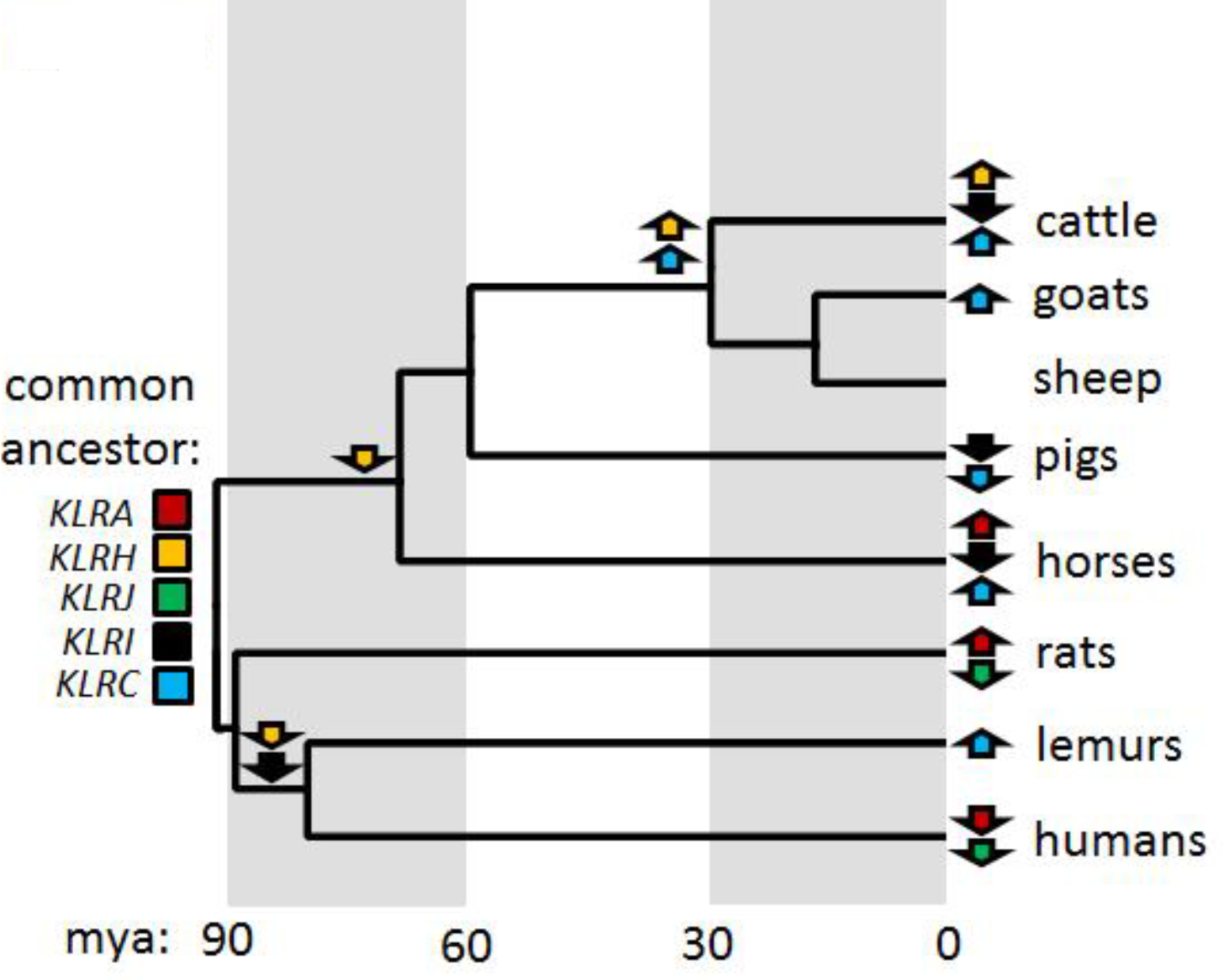
NKC evolution between selected mammalian lineages. Gene subgroup expansion (*up arrows*) or contraction (*down arrows*) is indicated at nodes. Divergence times are based on relative branch lengths and estimated using the reported divergence estimates for simians and prosimians (68.2-81.2 mya; Pozzi et al. (2014)), sheep and cattle (30 mya; Hiendleder et al. (1998)), and cattle and pigs (60 mya; (Meredith et al. 2011)). Alignment was generated using ClustalW (Thompson et al. 1994) and complete mitochondrial genomes (cattle, V00654; goat, KP271023; sheep, AF010406; pig, NC_000845; horse, AB859014; rat, X14848; mouse lemur, NC_028718; and human, AP008824). Phylogenetic trees were constructed using maximum likelihood based on the Tamura-3 parameter (Tamura 1992) model within MEGA6 (Tamura et al. 2013) using 100 bootstrap repetitions.

## DISCUSSION

### The evolution of the NKC

This study examined the NKC structure and gene content in eight mammalian species: cattle, sheep, goats, pigs, horses, rats, lemurs, and humans. This revealed extensive species-specific expansions and contractions amongst the various *KLR* gene subgroups within a generally conserved framework of genes. In particular, the *KLRA* locus has undergone extensive expansion and contraction during mammalian evolution, as evidenced by its highly variable gene content between species and the duplication and subsequent divergence of the closely related *KLRH* and *KLRJ* genes. All three of these genes originated prior to the divergence of the Laurasiatheria (e.g. cattle, pigs, whales, horses, dogs, etc.) and the Euarchontoglires (e.g. humans, lemurs, rabbits, rats, etc.) approximately 92 mya (Meredith et al. 2011). Within the human lineage, both *KLRH* and *KLRJ* were deleted (as well as *KLRI* and *KLRE* amongst all primates), and *KLRA* was rendered non-functional. Although different in rodents, *KLRH* is structurally similar in the mouse lemur and the Laurasiatherians, indicating that its sequence inverted and the intracellular and TM domains were deleted prior to the divergence of these two major clades. *KLRH* in rodents, on the other hand, is encoded in the same orientation as *KLRA* and possesses an inhibitory intracellular tail. This suggests either a duplication and subsequent contraction, or that a recombination occurred between the lectin domains of *KLRH* and a *KLRA* gene during rodent evolution. In either case, the inverted *KLRH*-like sequences were deleted in the rodent lineage. It seems likely that in early ruminant evolution this inverted region became a template for non-allelic homologous recombination (NAHR) in which sequence similarity between the regions containing *KLRC1* and *KLRH* established a conversion tract with *KLRC1* in the new location. The locus was then further expanded by duplication events and additional conversions. Thus, the duplication and expansion of the *KLRC* locus in ruminants would appear to be a consequence of the presence of the *KLRA*-related pseudogenes and fragments whose presence might create local instability driving evolutionary experimentation via conversion events with other *KLR* genes.

Phylogenetic analysis of the C-type lectin domains of the ruminant *KLRC* genes was unable to resolve the sequence of duplications, although it is likely to have initially occurred with an ancestor syntenic with *bota_KLRC1-7*. Shortly after the initial expansion, one of the now duplicated *KLRH* genes acquired the intracellular tail from an activating *KLRC2* gene. This new *KLRH*/*KLRC2* hybrid then subsequently duplicated along with the rest of the locus, giving rise to five such hybrids in cattle (*KLRH2*, *KLRH3*, *KLRH4*, *KLRH5*, and *KLRH6*) and one in goats (*cahi_KLRH2*). Three of these in cattle (*KLRH2*, *KLRH5* and *KLRH6*) are putatively functional. As the knockout mutations differ between the remaining three full-length pseudogenes, it is apparent that following the initial recombination event and subsequent expansion, these genes were independently disabled. This suggests that the *KLRH*/*KLRC2* hybrids were functional following the recombination event, and prior to their expansion. Intriguingly, the open reading frame for the C-type lectin domain of *KLRH* remains functionally preserved in cattle, sheep, goats, pigs, horses, and mouse lemurs, despite the lack of recognizable intracellular and TM domains. As it seems unlikely for this preservation to be coincidence, this gene may be functional, perhaps expressed as a soluble receptor.

The apparent conversion of activating receptors to pseudogenes in ruminants fits with a paradigm that activating genes are quickly expanded in response to strong selection pressure, then quickly lost once beneficial function is lost (Abi-Rached and Parham, 2005). In this hypothesis, retention of activating genes which are no longer useful to host survival and reproduction may be detrimental by permitting inappropriate NK cell stimulation, cytotoxicity, and autoimmunity. On the other hand, in the horse, all four identified pseudogenes contain inhibitory motifs. However, the three equine *KLRC1* subgroup pseudogenes appear to have undergone block duplication along with a functional *KLRC1*, thus expanding a net inhibitory receptor reservoir. The disruption of activating *KLRC2-1* in both cattle and goats appears to pre-date species divergence, as they share the same loss of the last two lectin domain exons. In cattle the loss of activating *KLRI2* function is either relatively recent or ongoing, due to the less obvious nature of the *knock* out mutations. That this gene is missing in pigs and horses suggests that it has either been lost, or its expansion has never occurred in their ancestry. The most parsimonius explanation is that they evolved once and subsequently homogenized within their respective species. Thus, as a result of gene conversion, the origin of many paired NK cell receptors is ambiguous.

### Use of long read sequencing to elucidate highly repetitive genomic regions

Much of the work described was motivated by the draft quality of the public genome assemblies for livestock species. Available methods using short-read sequencing data have difficulty forming high-quality assembly of repetitive areas of the genome such as the NKC. This difficulty is compounded when the two alleles present in the animal whose genome was sequenced may be substantially different in gene content in specific areas, since the assembly attempts to project a haploid presentation of the diploid genome. Some assemblers have tendency to collapse similar repeats on the assumption that they are allelic differences, while others expand the haploid genome to include more or all of the sequence from both alleles. In both instances, uncertainty usually results in a fractured assembly in the area of tandemly duplicated genes. This is simplified by sequencing haploid representations of the genome such as in large-insert clones in bacterial vectors. Indeed, the bovine and porcine genome assemblies, that included substantial content of sequence from individual BAC clones, provided significantly better (but still incomplete) representations of repetitive regions. But even with relatively short stretches of haploid DNA, short-read technologies have difficulty creating unbroken assemblies when the repeat length greatly exceeds the read length. Recently, long-read technologies have overcome many of the constraints of assembling these types of genomic regions (Koren et al. 2013; Berlin et al. 2015) and we employed these methods on some of the BAC clones used to obtain bovine genome sequence.

Our results also indicate the utility of using a probe-based capture method to enrich and sequence genomic regions to elucidate allelic variation. However, given the highly repetitive nature of the recently expanded *KLRC* genes in cattle, short-read paired-end (i.e. 250 bp × 250 bp) sequencing was insufficient for mapping and assessing the diversity across this region. Furthermore, the potential for genes containing domains that have recombined from other genes, such as the case for the cattle *KLRH/KLRC2* hybrids, may complicate efforts to transcriptomically assess gene expression when using standard short-read sequencing. To resolve these problems, we recommend that such repetitive immune gene clusters be sequenced and transcriptomically analyzed using long-read technology.

In conclusion, our annotations of the cattle, goat, sheep, pig, and horse NKC regions have identified a large proportion of NK cell receptor gene family members that may have been subjected to expansion and contraction due to NAHR. We described the extensive *KLRC* expansions in cattle and horses, the discovery of *KLRE* and *KLRI* outside of rodents, the presence of *KLRJ* in the described species, and the identification of novel *KLRH* genes bearing *KLRC2*-like activating tails in cattle and goats. Finally, polymorphisms across the cattle NKC, and likely other species, further expands the available NK cell receptor repertoire, particularly in the *KLRC/D* and *KLRA* systems. These results fill an important evolutionary link in our understanding of the NKC and will inform future investigations of NK cell receptor diversity, assist in identifying their potential ligands, and aid in the identification of genotypes associated with differential disease outcome.

## METHODS

### Ethics Statement

Peripheral blood samples from *B. taurus* and *B. indicus* cattle were collected in accordance with the U.K. Animal (Scientific Procedures) Act, 1986, and approved by either The Pirbright Institute Ethics Committee, or The Roslin Institute’s Animal Welfare and Ethics Committee. The Chillingham samples were from animals culled for welfare reasons. Blood sampling of *Kuchinoshima*-Ushi cattle was carried out in accordance with the Regulations for Animal Experiments in Nagoya University and the Guidelines for the Care and Use of Laboratory Animals by the Tokyo University of Agriculture.

### Genome assemblies

The region spanning the NKC from immediately upstream of *KLRA* to immediately downstream of *KLRE* was extracted from the current genome builds within Ensembl (Cunningham et al. 2015) for cattle (UMD_3.1, chr 5: 99387020-100235099), sheep (Oar_v3.1, chr 3:203826025-204418113), pig (Sscrofa10.2, chr 5:64112755-64634584), and horse (Equ Cab 2, chr 6:37209165-38556956), and from NCBI for goat (CHIR_1.0, CM001714: 91233093-91817092). Additional scaffolds for goat and cattle were generated using long-reads (NCBI accession numbers: PRJNA290100 and *KX592814*, respectively) and represent the first livestock genomes assembled *de novo* from PacBio reads alone. The specifics of sequence generation, contig assembly, scaffolding, and validation, to create the long-read assemblies is extensive and will be described elsewhere (Bickhart et al. 2016) (Smith and Medrano, unpublished). Genes within the NKC builds as well as on individual BAC clones were identified using the Basic Local Alignment Search Tool (BLAST) against GenBank and *known* NKC genes (Altschul et al. 1990). HMMgene was additionally used to hunt for putative open reading frames (Krogh 1997).

### Bacterial artificial chromosomes and sequencing

A BAC library was previously developed from a Friesian dairy bull (Di Palma 1999; Di Palma et al. 2002) and was screened for NKC-containing genes. Primers were designed to amplify *KLRC1*, *KLRD1*, *KLRD2*, *KLRJ*, and the flanking genes gamma-aminobutyric acid receptor-associated protein-like 1 (*GABARAPL1*) and serine/threonine/tyrosine kinase 1 (*STYK1*) (Supplementary Table 1). A PCR-based screen of 39,936 BAC clones (~1.5x genome coverage) identified three clones overlapping the NKC region: TPI4222-309A12, TPI4222-102F7, and TPI4222-343E2. Despite positive results with whole genomic DNA to verify primer specificity, however, no clones were PCR positive for *KLRJ* or *STYK1*. The NCBI genome viewer was queried to identify four additional BAC clones from two animals: two BAC clones from a different Friesian bull (RP42-154D6 and RP42-162P15), and two clones from the Hereford bull L1 Domino 99375 (CH240-60G5 and CH240-239G9).

BAC clones were expanded overnight and BAC DNA was purified using the Qiagen Large Construct *Kit* (Qiagen, GmbH). Purified DNA from three clones (TPI4222-309A12, TPI4222-102F7, and TPI4222-343E2) was sequenced using Illumina MiSeq with 250 bp × 250 bp paired-end reads (Source Biosciences, Inc.; Nottingham, United *Kingdom*) and *de novo* assembled using Velvet (Zerbino and Birney 2008). As the resultant assemblies failed to yield single contigs, the assembled sequences were manually scaffolded and supported by BLAST comparisons of individual contigs. These final, manual assemblies resulted in single contigs for each of the three clones. Assembly accuracy for these clones was confirmed by HindIII digestion of purified BAC DNA and comparison of band sizes to those predicted by assembly.

The remaining clones were sequenced at the USDA-ARS Meat Animal Research Center (Clay Center, Nebraska) using the PacBio RSII platform (Pacific Biosciences of California, Inc.). To further confirm the Illumina assemblies, TPI4222-309A12 and TPI4222-102F7 were re-sequenced in this manner as well. Read filtering and assembly was conducted using the Pacific Biosciences SMRT Analysis software (v2.3.0; http://www.pacb.com/devnet/). The resulting contigs were circularized by comparing the contig ends against the whole contigs to identify overlap, then the cloning vector identified and removed to produce contigs with the first base being the first beyond the 3’ end of the cloning vector accession sequence (AY487252). Potential errors remaining in the contig sequence were removed by re-mapping all of the subreads to the edited contigs, producing high quality (<0.01% error) genomic sequences.

### Nomenclature and manual annotation

Where possible, the Human Genome Organisation (HUGO) Gene Nomenclature Committee (HGNC)-approved gene nomenclature is used. For ease of reference, common gene synonyms are also provided upon first usage for many of the NKC genes. *KLRC* gene subgroup nomenclature is maintained based on the previous identification of *KLRC1* and *KLRC2*-like cDNA sequences in cattle (Birch and Ellis 2007). All NKC genome builds and individual BAC clones were manually annotated using Artemis (Rutherford et al. 2000). Pseudogenes were defined based on the presence of frameshifts and premature stop codons that would prevent the production of a functional protein. Recurrence plot sequence identity comparisons of genome assemblies were made using DOTTER (Sonnhammer and Durbin 1995) and a sliding window of 200 bp. Predictions of TM regions were made using TMHMM (Krogh et al. 2001). Alignments of NKC genes were generated using CLUSTALW (Thompson et al. 1994) and phylogenetic analyses were constructed within MEGA6 (Tamura et al. 2013) using maximum likelihood based on the Tamura three-parameter model and the partial deletion method using a 95% cut off (Tamura 1992).

### Animals used for SNP analysis

Heparinized peripheral blood was acquired from fifteen Friesian cattle (*B. taurus*) belonging to an MHC defined herd at The Pirbright Institute (Ellis et al. 1999). Semen from two Friesian breeding bulls (Blackisle Garve and Nerewater Tiptop) was purchased from Genus UK. Additional genomic DNA (gDNA) was obtained from two individuals from the feral Chillingham Park herd (Alnwick, Northumberland, United *Kingdom*) which have been genetically isolated for ~300 years (Visscher et al. 2001), an individual from a genetically isolated cattle population on *Kuchinoshima* Island (Japan) (Kawahara-Miki et al. 2011), two Sahiwal cattle (*B. indicus*) and a single Nelore (*B. indicus*). The gDNA sourced from each of the *B. taurus* and *B. indicus* animals were whole genome amplified using the REPLI-g mini kit (Qiagen, GmbH) following manufacturers’ instructions.

### Genomic enrichment of cattle NKC

Mononuclear cells (PBMCs) were separated from the peripheral blood of twenty *B. taurus* and three *B. indicus* cattle using Histopaque-1083 (Sigma-Aldrich Corporation) and gDNA was isolated using the QIAamp DNA mini kit (Qiagen, GmbH) following the manufacturers’ instructions. The quantity of purified gDNA was assessed for each animal with the Quant-iT PicoGreen dsDNA assay (Thermo Fisher Scientific) using the low range standard curve. An aliquot of gDNA from every animal was sheared using a Covaris S220 Focused-ultrasonicator (Covaris, Inc.). Instrument parameters provided by the manufacturer were used to fragment the DNA to insert sizes between 500-650 bp. Indexed paired-end gDNA libraries were constructed using a low-throughput, low-sample number TruSeq DNA sample preparation kit (Illumina, Inc.). Four multiplexed sequencing libraries were prepared, one for each of four independent genome enrichment experiments. Each multiplexed library was constructed using 1 μg input DNA per animal and ligation products were size selected (>500 bp) on an agarose gel and purified as described in the TruSeq protocol. An aliquot was removed from each and used as PCR template to assess the quality of the constructed sample library. PCR amplification was carried out as described in the TruSeq DNA Sample Preparation Guide. Sample library quality assessment was carried out by running PCR products on a DNA1000 chip using a 2100 Bioanalyzer instrument (Agilent Technologies).
Amplification of each multiplexed sample library was performed as described in the NimbleGen SeqCap EZ Library SR User’s Guide (v4.1). To enrich cattle gDNA from the NKC, custom oligonucleotide probes were designed and synthesised as a SeqCap EZ Developer Library (Roche Sequencing) and enrichment was performed as described in the manufacturers’ protocol. Human Cot-1 DNA was used to block repetitive regions of the cattle genome. DNA was purified at each stage of the Roche protocol using Agencourt AMPure XP DNA purification beads (Beckman Coulter, Inc.). The captured multiplex DNA sample was washed and recovered as outlined in the Roche protocol. Amplification of the enriched multiplex DNA sample libraries used LM-PCR as described in the Roche protocol. To determine how successful the enrichment was, DNA was analysed on a DNA1000 chip using a 2100 Bioanalyzer instrument (Agilent Technologies) and using qPCR. The degree of enrichment measured across the four independent captures indicated that each library was successfully generated.

### Sequencing and variant calling

The four enriched multiplex DNA sample libraries were independently sequenced using a MiSeq desktop sequencer (Illumina, Inc.) at the Pirbright Institute. The MiSeq Reagent *Kit* v2 (Illumina, Inc.) was used to produce either 2 × 230 bp or 2 × 250 bp paired-end reads per run. Multiplexed DNA sample libraries were diluted and sequenced at a final concentration of 8 pM. Each sequencing run used a PhiX control spike which was denatured and diluted to 12.5 pM. The final pool of sequenced DNA was comprised of 99% sample library and 1% PhiX. Library preparation, sample loading, and MiSeq preparation steps were carried out as described in the manufacturers’ protocol. Resultant reads were mapped to the cattle genome build UMD_3.1 using the Burrows-Wheeler Aligner (BWA, v0.7.5a) (Li and Durbin 2009) and variant sites were identified using SAMtools (v0.1.18) (Li et al. 2009) and VarScan (v2.3.6) (Koboldt et al. 2012).

## DATA ACCESS

The long read genome scaffolds for the goat assembly (ARS1; BioProject: PRJNA290100 and assembly: LWLT00000000) and the 12.7 mb scaffold containing the NKC from the long-read assembly of the cattle genome (GenBank: KX592814) have been deposited with NCBI. The GenBank accessions for the BAC clones used in the present study are: TPI4222-343E2 (KX611578), TPI4222-309A12 (KX611577), TPI4222-102F7 (KX611576), RP42-162P15 (KX698608), RP42-154D6 (KX698607), CH240-239G9 (AC170009), and CH240-60G5 (AC156849).

## ACKNOWLEDGEMENTS

We thank William Thompson for excellent technical support. JCS and JAH were supported by the United Kingdom Biotechnology and Biological Sciences Research Council (BBSRC) Institute Strategic Program on Livestock Viral Diseases awarded to The Pirbright Institute. MSG was supported by BBSRC grant BB/J006211/1, “Dissecting the functional impact of natural killer cell receptor variation in cattle.” DMB was supported in part by appropriated project 1265-31000-096-00, “Improving Genetic Predictions in Dairy Animals Using Phenotypic and Genomic Information”, of the Agricultural Research Service of the United States Department of Agriculture. DMB and TPLS were also supported by the Agricultural Food Research Initiative (AFRI) competitive grant number: 2015-67015-22970 from the USDA National Institute of Food and Agriculture (NIFA) Animal Health Program. We would like to thank Prof Elizabeth Glass (The Roslin Institute, UK) for providing the Sahiwal and Nelore DNA samples, Prof Tomohiro Kono (Tokyo University of Agriculture, Japan) for providing the Kuchinoshima-Ushi DNA, and Prof Stephen Hall (University of Lincoln, UK) for useful comments and for providing peripheral blood from the culled Chillingham cattle. Mention of trade names or commercial products in this article is solely for the purpose of providing specific information and does not imply recommendation or endorsement by the US Department of Agriculture.

